# CRISPR-Cas12a genome editing at the whole-plant level using two compatible RNA virus vectors

**DOI:** 10.1101/2021.04.19.440450

**Authors:** Mireia Uranga, Marta Vazquez-Vilar, Diego Orzáez, José-Antonio Daròs

## Abstract

The use of viral vectors that can replicate and move systemically through the host plant to deliver bacterial clustered, regularly interspaced, short palindromic repeats (CRISPR) components enables genome editing at the whole-plant level and avoids the requirement for labor-intensive stable transformation. However, this approach usually relies on previously transformed plants that stably express a CRISPR-associated (Cas) nuclease. Here we describe successful DNA-free genome editing of *Nicotiana benthamiana* using two compatible RNA virus vectors, derived from tobacco etch virus (TEV; genus *Potyvirus*) and potato virus X (PVX; genus *Potexvirus*), which replicate in the same cells. The TEV and PVX vectors respectively express a Cas12a nuclease and the corresponding guide RNA. This novel two-virus vector system improves the toolbox for transformation-free virus-induced genome editing in plants and will advance efforts to breed more nutritious, resistant, and productive crops.

## Introduction

Systems derived from bacterial clustered, regularly interspaced, short palindromic repeats (CRISPR) and CRISPR-associated (Cas) proteins (Cong *et al*., 2013) have revolutionized biotechnology. In plants, CRISPR-Cas holds great promise for unprecedented genome engineering of both model species and crops (Nekrasov *et al*., 2013; Zhang *et al*., 2016; Li *et al*., 2017; Zhu *et al*., 2020; Huang and Puchta, 2021). Most common CRISPR/Cas arrangements include a Cas endonuclease, such as *Streptococcus pyogenes* SpCas9, and a single-guide RNA (sgRNA), which specifically directs the nuclease to a sequence of interest in the genome. As in other taxonomic groups (Platt et al., 2014; Senís et al., 2014; Lau and Suh, 2017; Xu et al., 2019), virus-derived vectors have been reported as a powerful alternative to express the CRISPR/Cas components at the whole-plant level avoiding the labor-intensive and time-consuming tissue culture approaches required for stable transformation. These strategies are commonly termed as virus-induced genome editing (VIGE) and have focused on the delivery of one or more sgRNAs using RNA or DNA virus vectors in transgenic plants that stably express the Cas nuclease (Ali *et al*., 2015; Yin *et al*., 2015; Cody *et al*., 2017; Ali *et al*., 2018; Hu *et al*., 2019; Jiang *et al*., 2019; Ellison *et al*., 2020; Lei *et al*., 2021).

Expression of a Cas nuclease using a plant virus-derived vector able to move systemically through a plant was long considered unachievable due to cargo constraints. However, the innovative work by Ma et al. (2020) demonstrated efficient genome editing by delivering both the sgRNA and SpCas9 at the whole-plant level using a vector derived from sonchus yellow net virus (SYNV; family *Rhabdoviridae*). Despite this unprecedented achievement, additional virus-based systems for the co-expression of Cas nucleases and sgRNAs at the whole-plant level are still required to improve the current toolbox for crop engineering. Notably, each viral vector has its own unique properties, particularly a specific host range.

To expand the virus-based tools for tissue culture-free genome editing in plants, we co-expressed the Cas nuclease and the guide RNA using two compatible viral vectors that replicate in the same cells and coordinately move systemically through the whole-plant. We chose a potyvirus vector to express the Cas nuclease. Potyviruses (genus *Potyvirus*) are the largest group of plus-strand RNA viruses, with more than 200 currently known species that collectively can infect a large range of host plants (Wylie *et al*., 2017). We also focused on a Cas12a (formerly Cpf1) nuclease, which is a component of a class 2 type V CRISPR system, isolated from *Lachnospiraceae bacterium* ND2006 (LbCas12a) (Zetsche et al., 2015). Genome editing using LbCas12a has been demonstrated in plants (Endo et al., 2016; Tang et al., 2017; Wang et al., 2017; Xu et al., 2017), and the complementary DNA (cDNA) corresponding to this nuclease is smaller than that of SpCas9. We also co-expressed the guide RNA using a recently described potato virus X (PVX, genus *Potexvirus*, family *Virgaviridae*) vector, which efficiently induces hereditable gene editing in *Nicotiana benthamiana* plants that stably express SpCas9 (Uranga *et al*., 2021). Our results demonstrated efficient DNA-free *N. benthamiana* genome editing using the two compatible RNA virus vectors.

## Methods

### Viral vectors

Guide RNAs to target *N. benthamiana Flowering locus T* (*NbFT*; SolGenomics Niben101Scf01519g10008.1) and *Xylosyl transferase 1* (*NbXT1*; Niben101Scf04205g03008.1) were selected using the CRISPR-P online tool as described by Bernabé-Orts et al. (2019) (Table S1). Nucleotide (nt) sequences of recombinant viral clones are shown in Figs. S1 and S2. These clones were built using the primers shown in Tables S2 and S3.

### Plant inoculation

*N. benthamiana* wild-type and transformed plants expressing tobacco etch virus (TEV, genus *Potyvirus*) nuclear inclusion *b* (NIb) protein (Martí *et al*., 2020), were grown at 25°C under a 12 h/12 h day/night photoperiod. Plants that were 4-to 6-weeks-old were agroinoculated as previously described (Bedoya *et al*., 2010; Uranga *et al*., 2021). Tissue samples (approximately 100 mg) from the first symptomatic upper non-inoculated leaf were collected at different days post inoculation (dpi), as indicated, for virus progeny and plant genome-editing analyses.

### Reverse transcription (RT)-polymerase chain reaction (PCR) analysis of viral progeny

RNA was purified from leaf samples using silica gel columns (Uranga *et al*., 2021). cDNA was synthesized using RevertAid reverse transcriptase (Thermo Scientific) and primer D179 (Table S4). PCR with *Thermus thermophilus* DNA polymerase (Biotools) was used to amplify the TEV coat protein (CP) cistron (primers D178 and D211; Table S4) or a fragment of the LbCas12a open reading frame (ORF) (primers D3604 and D3605; Table S4). PCR products were separated by electrophoresis in 1% agarose gels followed by staining with ethidium bromide.

### Analysis of *N. benthamiana* genome editing

DNA from leaf samples was purified using silica gel columns (Uranga *et al*., 2021). *N. benthamiana* genome fragments were amplified by PCR using high-fidelity Phusion DNA polymerase (Thermo Scientific) (Table S5). PCR products were separated by agarose gel electrophoresis, purified from the gel, and subjected to Sanger sequencing (Table S5). The presence of sequence modifications was analyzed using the inference of CRISPR edits (ICE) software (http://www.synthego.com/products/bioinformatics/crispr-analysis).

## Results

### A dual virus-based vector system to co-express Cas nucleases and guide RNAs in plants

Initially, we built a TEV recombinant clone in which the cDNA of a human codon-optimized LbCas12a replaced that of the viral NIb protein (TEVΔN::LbCas12a). Our previous work showed that this vector could express large exogenous sequences, such as a whole bacterial metabolic pathway to biosynthesize lycopene or the saffron carotenoid cleavage dioxygenase (Majer *et al*., 2017; Martí *et al*., 2020). However, since the virus lacks the viral RNA-dependent RNA polymerase NIb, it replicates only in plants that express this protein (Majer *et al*., 2017). In our recombinant clone, the sequence coding for LbCas12a replaced most of the NIb cistron and was flanked by the native nuclear inclusion *a* protease (NIaPro) cleavage sites that mediate the release of the nuclease from the viral polyprotein (Fig. 1A and Fig. S1).

**FIG. 1.**
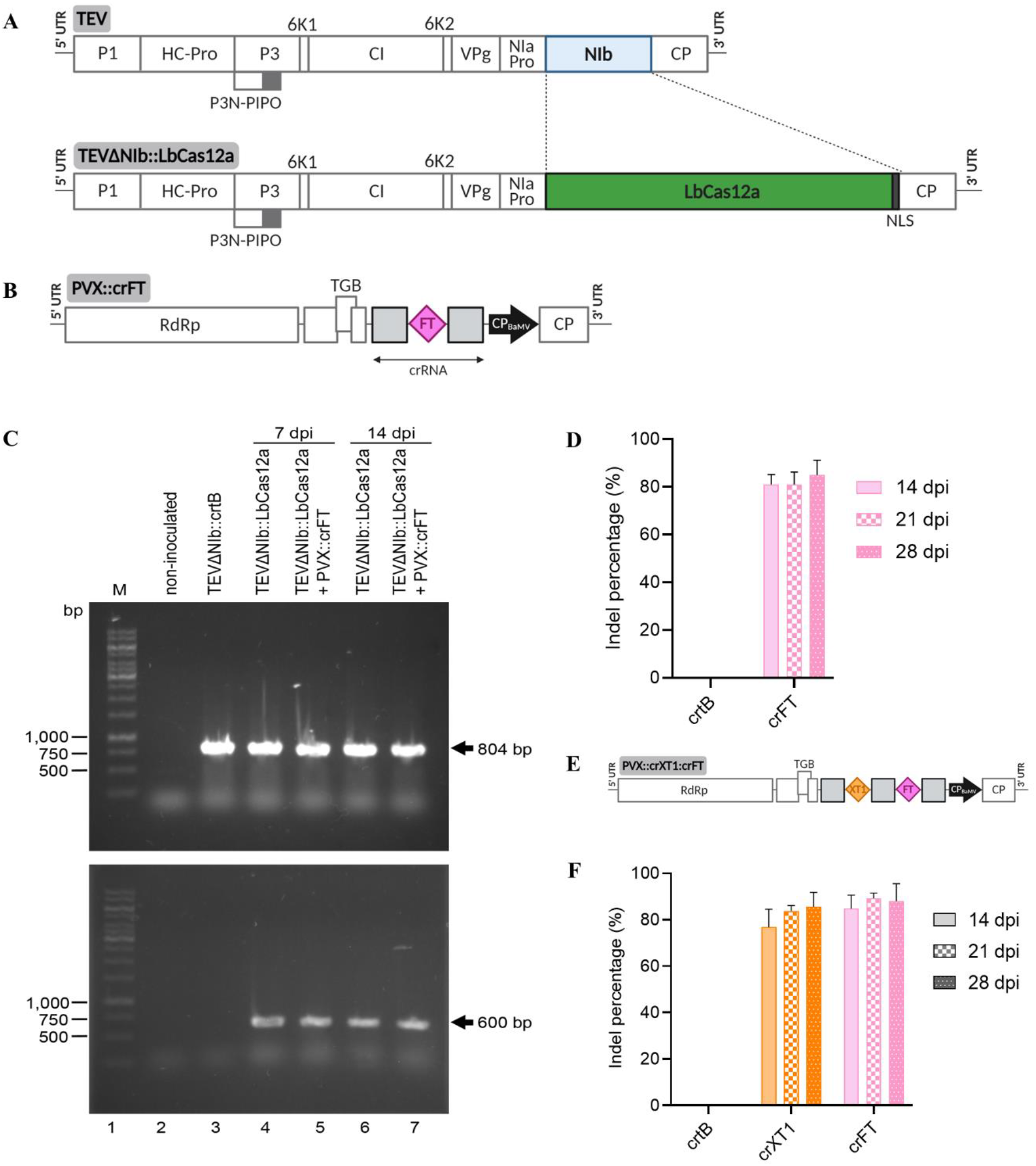
DNA-free gene editing in *N. benthamiana* based on virally delivered LbCas12a and crRNA. **(A)** Schematic representation of recombinant virus TEVΔNIb::LbCas12a. Cas12a ORF from *Lachnospiraceae bacterium* ND2006 (LbCas12a), containing a carboxy-terminal nucleoplasmin nuclear localization signal (NLS, grey box), replaces the TEV NIb cistron (light blue box) and is flanked by proteolytic processing sites of viral NIaPro. TEV cistrons P1, HC-Pro, P3, P3N-PIPO, 6K1, CI, 6K2, VPg, NIaPro, NIb, and CP are represented by boxes. 5’ and 3’ untranslated regions (UTRs) are represented by lines. **(B)** Schematic representation of recombinant clone PVX::crFT. RNA-dependent RNA polymerase (RdRp), triple gene block (TGB), and coat protein (CP) are represented by open boxes. Heterologous *Bamboo mosaic virus* CP promoter (CP_BaMV_) is represented by a black arrow. 5’ and 3’ UTRs are represented by black lines. crRNA consists of a gene-specific 23-nt protospacer (pink diamond) and conserved 21-nt direct repeats (grey boxes). CP_BaMV_, protospacer and scaffold are not shown at scale. **(C)** RT-PCR analysis of TEVΔNIb progeny at 7 and 14 dpi in 35S::NIb *N. benthamiana* plants inoculated with TEVΔNIb::LbCas12a alone or co-inoculated with TEVΔNIb::LbCas12a and PVX::crFT. Amplification products were separated by electrophoresis in an agarose gel stained with ethidium bromide. Lane 1, 1-kb ladder DNA marker with the length of some components (in bp) indicated on the left; lane 2, non-inoculated plant; lanes 3 to 7, plants inoculated with TEVΔNIb::crtB (lane 3) or TEVΔNIb::LbCas12a (lanes 4 and 6), and co-inoculated with TEVΔNIb::LbCas12a and PVX::crFT (lanes 5 and 7). The amplification products corresponding to cDNA regions of TEV CP (804 bp) (top) or LbCas12a (600 bp) (bottom) are indicated by arrows. **(D)** ICE analysis of the first systemically infected upper leaf of *N. benthamiana* plants co-inoculated with TEVΔNIb::LbCas12a and PVX::crFT at the indicated dpi (n=6). TEVΔNIb::crtB was used as a negative control. (**E**) Schematic representation of recombinant clone PVX::crXT1:crFT. Protospacers for NbXT1 and NbFT are represented by orange and pink diamonds, respectively. Other details are indicated above. (**F**) ICE analysis of the first systemically infected upper leaf of *N. benthamiana* plants (n=6) co-inoculated with TEVΔNIb::LbCas12a and PVX::crXT1:crFT at the indicated dpi. Columns and error bars represent average indels (%) and standard deviation, respectively.

Next, we built a recombinant version of PVX in which the LbCas12a CRISPR RNA (crRNA) was expressed under the control of the viral CP promoter, and the 29 initial codons of PVX CP were deleted to improve the stability of the recombinant clone (Dickmeis et al., 2014). The viral CP was expressed from a heterologous promoter derived from that of the CP of bamboo mosaic virus (BaMV; genus *Potexvirus*). Based on previous work by Bernabé-Orts et al. (2019) assessing the efficiency of the Cas12a-mediated gene editing of several *N. benthamiana* loci, *NbFT* was selected as the target gene (PVX::crFT) (Fig. 1B and Fig. S2). The 65-nt crRNA was cloned downstream of the PVX CP promoter and consisted of a 23-nt protospacer sequence specific to the target gene flanked on both 5’ and 3’ ends by a conserved 21-nt scaffold, also known as a direct repeat.

*N. benthamiana* plants constitutively expressing TEV NIb under the control of cauliflower mosaic virus (CaMV) 35S promoter and terminator (Martí *et al*., 2020) were co-inoculated with a 1:1 mix of two cultures of *Agrobacterium tumefaciens* transformed with plasmids harboring TEVΔNIb::LbCas12a and PVX::crFT constructs. Controls included inoculation of TEVΔNIb::LbCas12a alone and inoculation of TEVΔNIb::crtB, which allows visual tracking of virus systemic movement due to the yellow pigmentation of infected tissue induced by *Pantoea ananatis* phytoene synthase (crtB) (Majer et al., 2017). At 7 dpi, typical symptoms of TEV infection emerged in the upper non-inoculated leaves of all plants. Notably, at 14 dpi irregular chlorotic spots appeared in the leaves of plants co-inoculated with TEVΔNIb::LbCas12a and PVX::crFT, but not in those inoculated with TEVΔNIb::LbCas12a alone (Fig. S3). PVX infection is characterized by the appearance of vein banding, ring spots and leaf atrophy (Loebenstein and Gaba, 2012). Therefore, the observed phenotypic alteration could have been due to LbCas12a-mediated editing of *NbFT* or an effect of the coinfection. Samples from the first systemically infected upper leaf were collected at 7 and 14 dpi, and TEVΔNIb progeny were studied by RT-PCR analysis (Fig. 1C). An 804-bp specific region of the TEV genome corresponding to the CP cistron was amplified in all virus-inoculated plants, confirming the presence of the virus (Fig. 1C, top panel). However, for viral vectors carrying big cargos, genomes larger than wild type are likely to recombine to smaller sizes, thus triggering the loss of heterologous genes (Gilbertson et al., 2003). An additional RT-PCR analysis was performed, and a 600-bp cDNA corresponding to a fragment of the LbCas12a ORF was exclusively amplified from plants inoculated with TEVΔNIb::LbCas12a alone or co-inoculated with TEVΔNIb::LbCas12a and PVX::crFT, regardless of sampling time (Fig. 1C, bottom panel). These results suggested that the LbCas12a nuclease was expressed from the onset of viral infection and throughout the rest of the experiment.

Next, DNA was purified from leaf samples collected at 7 and 14 dpi, and a 550-bp fragment of the *NbFT* gene covering the LbCas12a target site was amplified by PCR. Sanger sequencing of the PCR products and ICE analysis revealed robust gene editing at 14 dpi in plants co-inoculated with TEVΔNIb::LbCas12a and PVX::crFT, reaching an indel percentage of up to 75% (Fig. 1D and Fig. S4). No significant differences were observed at 21 and 28 dpi. The indel distribution was consistent with the deletion-enriched mutagenesis profile characteristic of Cas12a activity, being mainly 5-to 10-bp deletions (Bernabé-Orts et al., 2019). These results indicate that the simultaneous delivery of CRISPR-Cas12a components through two compatible viral vectors (i.e., TEVΔNIb and PVX) allows highly efficient, DNA-free targeted mutagenesis in *N. benthamiana*.

### Multiplex genome editing using the dual virus-based vector system in plants

A key advantage of CRISPR-Cas genome editing is the capacity to target several loci at once by the simultaneous expression of several guide RNAs (i.e., multiplexing). Based on our observations in Cas9-expressing plants (Uranga *et al*., 2021), we wondered whether the PVX vector could allow the delivery of multiple, functional crRNAs for Cas12a-mediated editing. To investigate this, we selected *Xylosyl transferase 1* (*NbXT1*) as the second target gene. As Cas12a can self-process crRNAs due to its RNase III activity, *NbXT1* and *NbFT* crRNAs were arranged in tandem under the control of the same CP promoter, thus creating the PVX::crXT1:crFT construct (Fig. 1E). *N. benthamiana* plants constitutively expressing TEV NIb were co-inoculated with a 1:1 mix of two *A. tumefaciens* cultures carrying TEVΔNIb::LbCas12a and PVX::crXT1:crFT. *A. tumefaciens* transformed with TEV::crtB was used as a control in this assay. The first systemically infected leaf was sampled at 14, 21, and 28 dpi, following extraction of genomic DNA and PCR amplification of the target sites. ICE analysis revealed efficient gene editing on both *NbXT1* and *NbFT*, with average indel percentages ranging from 76% to 88%, which was maintained regardless of the sampling time (Fig. 1F). In addition, the absence of statistically relevant differences in gene editing among *NbXT1* and *NbFT* suggested that LbCas12a can efficiently self-process tandemly arrayed crRNAs. Time-course comparison with the single-crRNA construct, in the case of *NbFT*, also revealed that multiplexing does not affect the editing efficiency, since the mutation rates achieved with both strategies were similar (Fig. 1D and F).

### Dual vector CRISPR-Cas genome editing in wild-type plants

The fact that TEVΔNIb infectivity depends on the supplementation of viral NIb from a transgene may be perceived as a limitation of this genome-editing system, as the approach is still bound to a previously transformed plant. An alternative strategy for supplying NIb activity consists of the co-inoculation of TEVΔNIb with a recombinant PVX expressing NIb (Bedoya et al., 2010). We wondered whether a single PVX vector could: (i) provide NIb activity for the systemic movement of TEVΔNIb; and (ii) deliver the crRNA for LbCas12a-mediated gene editing. Thus, the coding sequence of the TEV NIb cistron plus an additional amino-terminal Met was inserted within the expression cassette in PVX. *NbFT*-specific crRNA was added downstream of NIb without any linker sequence, so that both NIb and crFT expression were under the control of the PVX CP promoter (PVX::NIb:crFT) (Fig. 2A). Wild-type *N. benthamiana* plants were co-inoculated with a 1:1 mix of two cultures of *A. tumefaciens* carrying TEVΔNIb::LbCas12a and PVX::NIb:crFT. *A. tumefaciens* transformed with TEV::crtB was again used as a control. At 7 dpi, the apical leaves of the co-inoculated *N. benthamiana* plants became symptomatic (i.e., leaf curling). At 14 dpi, necrotic spotting and interveinal mottling were observed in systemic leaves, which became more noticeable over time (Fig. S5). Samples from the first systemically infected upper leaf were collected at 14 dpi and the 550-bp fragment of the *NbFT* gene covering the LbCas12a target site was amplified by PCR. ICE analysis of the PCR products exhibited 20% indels in plants co-inoculated with TEVΔNIb::LbCas12a and PVX::NIb:crFT. These results indicate that a single PVX vector can supply the viral NIb activity that allows TEVΔNIb to systemically spread in wild-type *N. benthamiana*, as well as to perform crRNA delivery for LbCas12a-mediated genome editing.

**FIG. 2.**
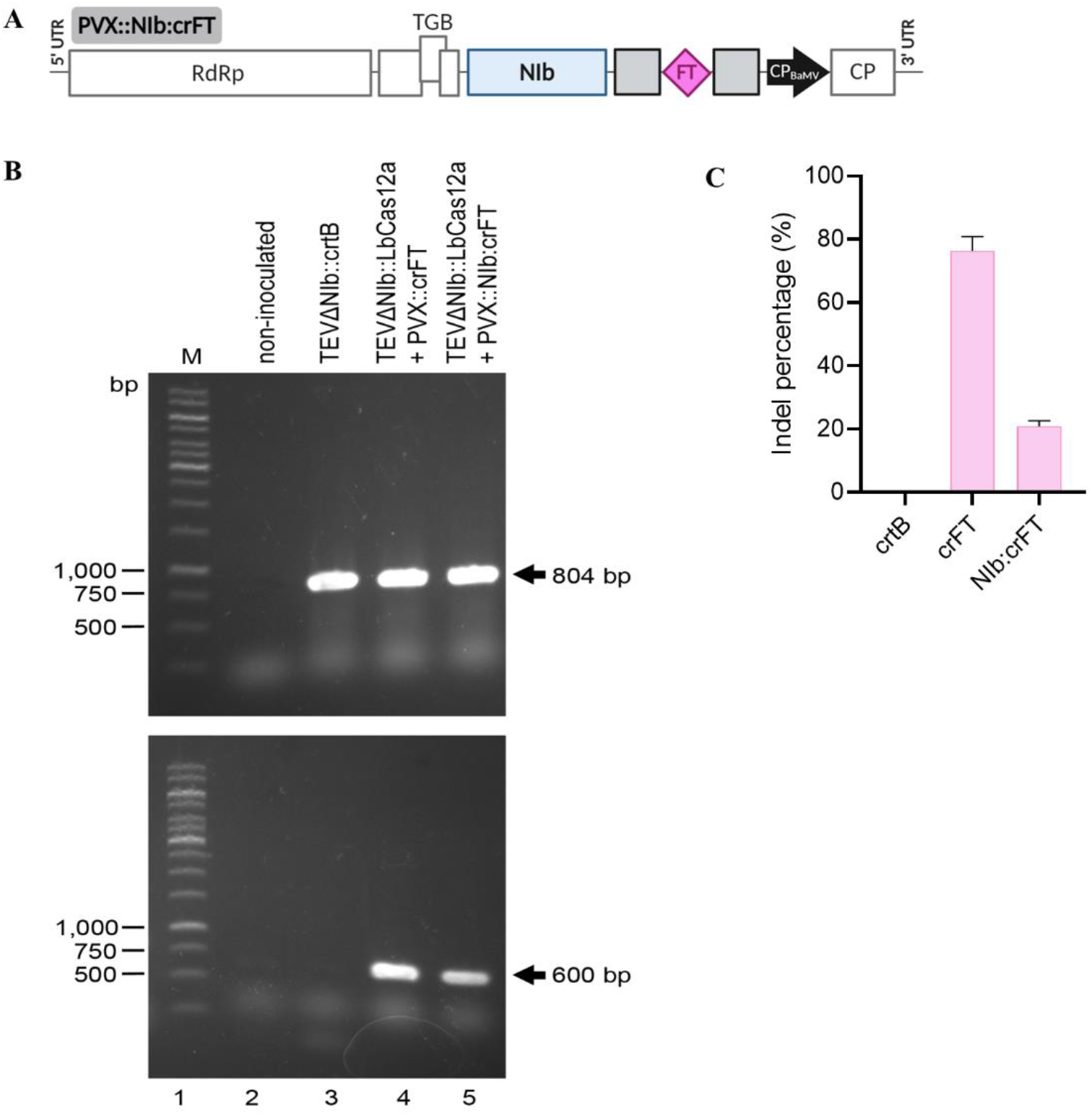
Engineering of a single PVX vector for complementation of defective TEVΔNIb and expression of LbCas12a crRNA. **(A)** Schematic representation of recombinant virus PVX::NIb:crFT. TEV NIb cistron is represented by a light blue box. Other details are as described in the legend to Fig. 1. **(B)** RT-PCR analysis of TEVΔNIb progeny at 14 dpi in wild-type *N. benthamiana* plants co-inoculated with TEVΔNIb::LbCas12a and PVX::crFT or with TEVΔNIb::LbCas12a and PVX::NIb:crFT. Amplification products were separated by electrophoresis in an agarose gel that was stained with ethidium bromide. Lane 1, DNA marker ladder with the length of some components (in bp) indicated on the left; lane 2, non-inoculated plant; lane 3 to 5, plants inoculated with TEVΔNIb::crtB (lane 3) and co-inoculated with TEVΔNIb::LbCas12a and PVX::crFT (lane 4) or with TEVΔNIb::LbCas12a and PVX::NIb:crFT (lane 5). The amplification products corresponding to cDNA regions of TEV CP (804 bp) (top) or LbCas12a (600 bp) (bottom) are indicated by arrows. (c) ICE analysis of the first systemically infected upper leaf of *N. benthamiana* plants co-inoculated with TEVΔNIb::LbCas12a and PVX::NIb:crFT at 14 dpi (n=6). TEVΔNIb::crtB was used as a negative control. Columns and error bars represent average indels (%) and standard deviation, respectively.

## Discussion

In this study we describe the engineering of a dual RNA-virus system for the delivery of CRISPR-Cas12a components for genome editing in plants. The system consists of two compatible plus-strand RNA viruses, specifically the potyvirus TEV and the potexvirus PVX, to express the Cas nuclease and the guide RNA, respectively. Notably, the TEV vector (TEVΔNIb) contains the deletion of the RNA-dependent RNA polymerase (NIb) in order to accommodate the large ORF of the Cas nuclease. Our previous efforts to express a functional Cas nuclease using a full-length potyvirus vector in plants have been unsuccessful. However, recent works have reported successful Cas9 expression and DNA-free genome editing using single SYMV (Ma et al., 2020) and PVX (Ariga *et al*., 2020) vectors in *N. benthamiana*. These contrasting results emphasize the diverse properties of vectors derived from viruses belonging to different genera and families, and the necessity for a large spectrum of molecular tools, operating in a wide range of host species, to tackle challenging VIGE goals in crop plants. Notably, SYNV is a minus-strand RNA virus the inoculation of which entails some complexity (Peng *et al*., 2021) and, as with each plant virus, it exhibits a particular host range (Jackson and Christie, 1977).

In this work, we aimed to express a Cas nuclease using a potyvirus vector for DNA-free plant-genome editing. The genus *Potyvirus* is the largest among the plant RNA viruses, and cromprises more than 200 species that infect a wide range of host plants from many different botanical families (Wylie *et al*., 2017). Therefore, it offers a wealth of genetic resources for VIGE. Although we were unable to successfully express the Cas nuclease using a full-length potyvirus vector, we demonstrated that TEVΔNIb allows the transient expression of LbCas12a, and that PVX can perform both single or multiple crRNA delivery as well as providing the NIb activity. Our two-virus delivery system resulted in efficient DNA-free targeted editing both in NIb-expressing and wild-type *N. benthamiana* plants, reaching indel percentages of up to 80% and 20%, respectively (Figs. 1 and 2). This dual vector system not only incorporates the enormous genetic resources of potyviruses for VIGE, but also demonstrates that compatible RNA virus vectors can be used to simultaneously deliver several CRISPR-Cas components. This could be useful for more sophisticated arrangements in DNA-free plant-genome editing and gene-expression regulation studies. Recently, we developed a PVX vector to efficiently express multiplex SpCas9 sgRNAs at the whole-plant level. Current results with LbCas12a confirm that PVX can be easily engineered for the simultaneous delivery of guide RNAs regardless of the nature of the Cas nuclease, which highlights its usefulness in a variety of multiplexing approaches.

In contrast to previous reports of DNA-free VIGE that used SpCas9 (Ariga *et al*., 2020; Ma *et al*., 2020), we selected Cas12a. This was due to the following unique features of the nuclease (Zaidi et al., 2017): (i) the cleavage of target DNA is directed by a single crRNA shorter than that of SpCas9 sgRNA; (ii) the protospacer adjacent motif (PAM) is T-rich (5’-TTTN-3’); (iii) DNA cleavage results in cohesive ends with 4-or 5-nt overhangs, which might facilitate homology-directed repair (HDR); (iv) it exhibits RNase III activity useful to facilitate multiplex gene editing; and (v) it is smaller than SpCas9 (3.8 kb vs 4.2 kb), which is important in terms of viral delivery. Moreover, the unique characteristics of the CRISPR-Cas12a system, unlike that of Cas9, such as the recognition of a T-rich PAM and the induction of staggered ends that facilitate homologous recombination, limit the range of target sequences and thus reduce off-target activity (Zaidi et al., 2017). Cas12a orthologs from *Francisella novicida* U112 (FnCas12a), *Acidaminococcus* sp. BV3L6 (AsCas12a) and LbCas12a were first experimentally validated in mammalian cells (Kim et al., 2016, 2017a; Zetsche et al., 2017). In plants, targeted mutagenesis was achieved in rice and tobacco using any of the orthologs (Endo et al., 2016; Tang et al., 2017; Wang et al., 2017; Xu et al., 2017). A DNA-free approach based on the delivery of AsCas12a or LbCas12a loaded with crRNA was also validated in wild tobacco and soy-bean protoplasts (Kim et al., 2017b). Here we focused on LbCas12a, since previous works reported that this nuclease possesses higher efficiency than FnCas12a or AsCas12a (Tang et al., 2017; Bernabé-Orts et al., 2019), and is also effective for plant-genome editing when virally delivered.

In conclusion, our dual RNA-virus-based system broadens the current toolbox for DNA-free VIGE and will contribute to applications in plant functional genomics and crop improvement.

## Supplementary Material

Figure S1. Full sequence of wild-type tobacco etch virus (TEV, Genbank accession number DQ986288), the defective viral vector TEVΔNIb and their derived recombinant viruses, TEV::crtB, TEVΔNIb::crtB and TEVΔNIb::LbCas12a.

Figure S2. Full sequence of wild-type potato virus X (PVX; GenBank accession number MT799816) and its derived recombinant viruses PVX::crFT and PVX::NIb:crFT.

Figure S3. 35S::NIb *N. benthamiana* plants constitutively expressing TEV NIb and representative leaves from these plants at 14 dpi inoculated with TEVΔNIb::LbCas12a or co-inoculated with TEVΔNIb::LbCas12a and PVX::crFT.

Figure S4. Example of a sequence electropherogram and mutagenesis profile from 35S::NIb N. benthamiana plants co-inoculated with TEVΔNIb::LbCas12a and PVX::crFT.

Figure S5. Representative leaves from wild-type *N. benthamiana* plants at 14 dpi inoculated with TEVΔNIb::LbCas12a alone, PVX::NIb:crFT alone, or coinoculated with TEVΔNIb::LbCas12a and PVX::NIb:crFT.

Table S1. *N. benthamiana* genes targeted by the CRISPR-Cas12a system. PAMs are highlighted with a grey background.

Table S2. Primers used for the construction of recombinant viruses.

Table S3. Primer combinations used for the construction of recombinant viruses. Table S4. Primers used for TEVΔNIb diagnosis by RT-PCR.

Table S5. Primers used for Cas12a-crRNA gene editing analysis.

## SUPPLEMENTAL DATA

**Figure S1.**
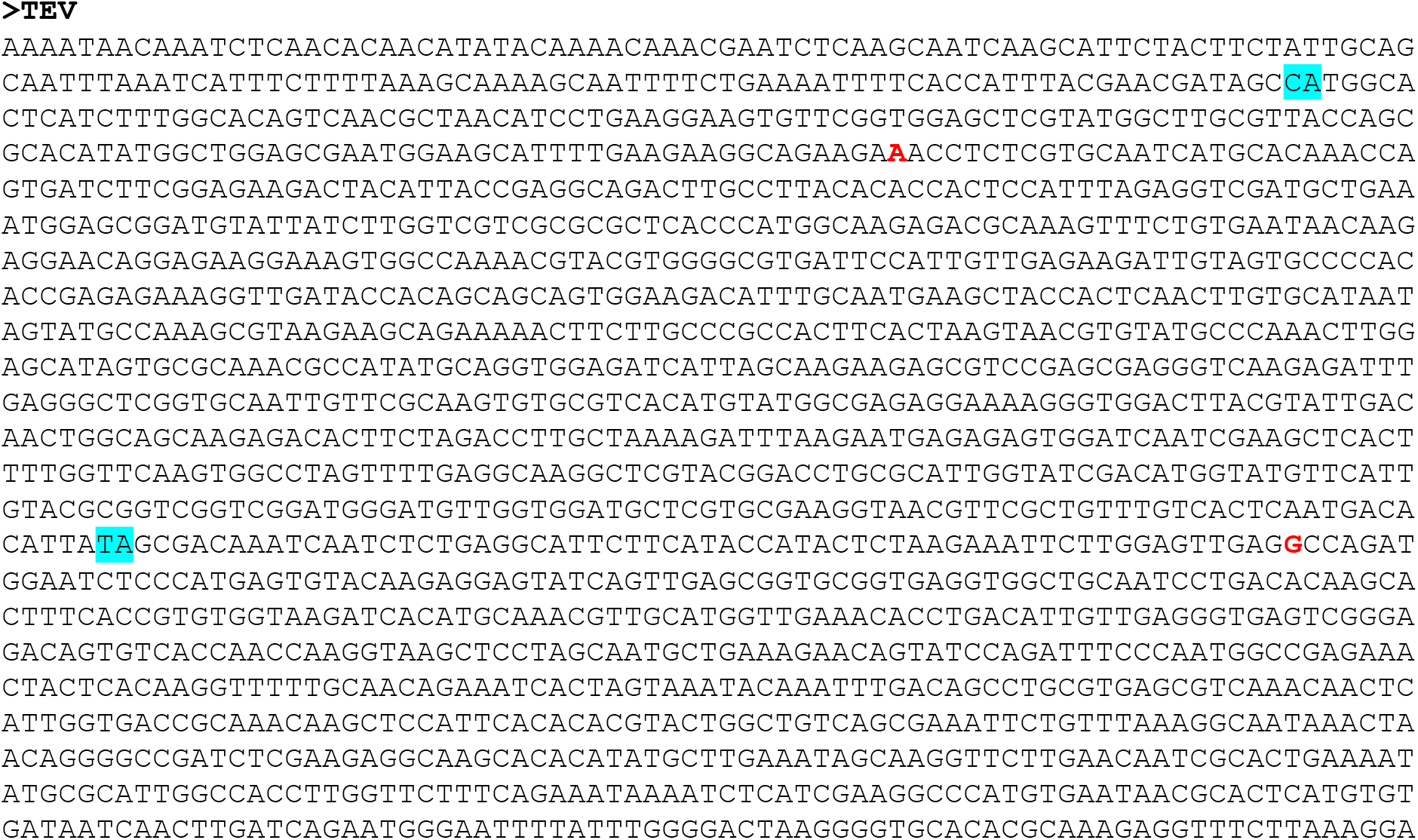
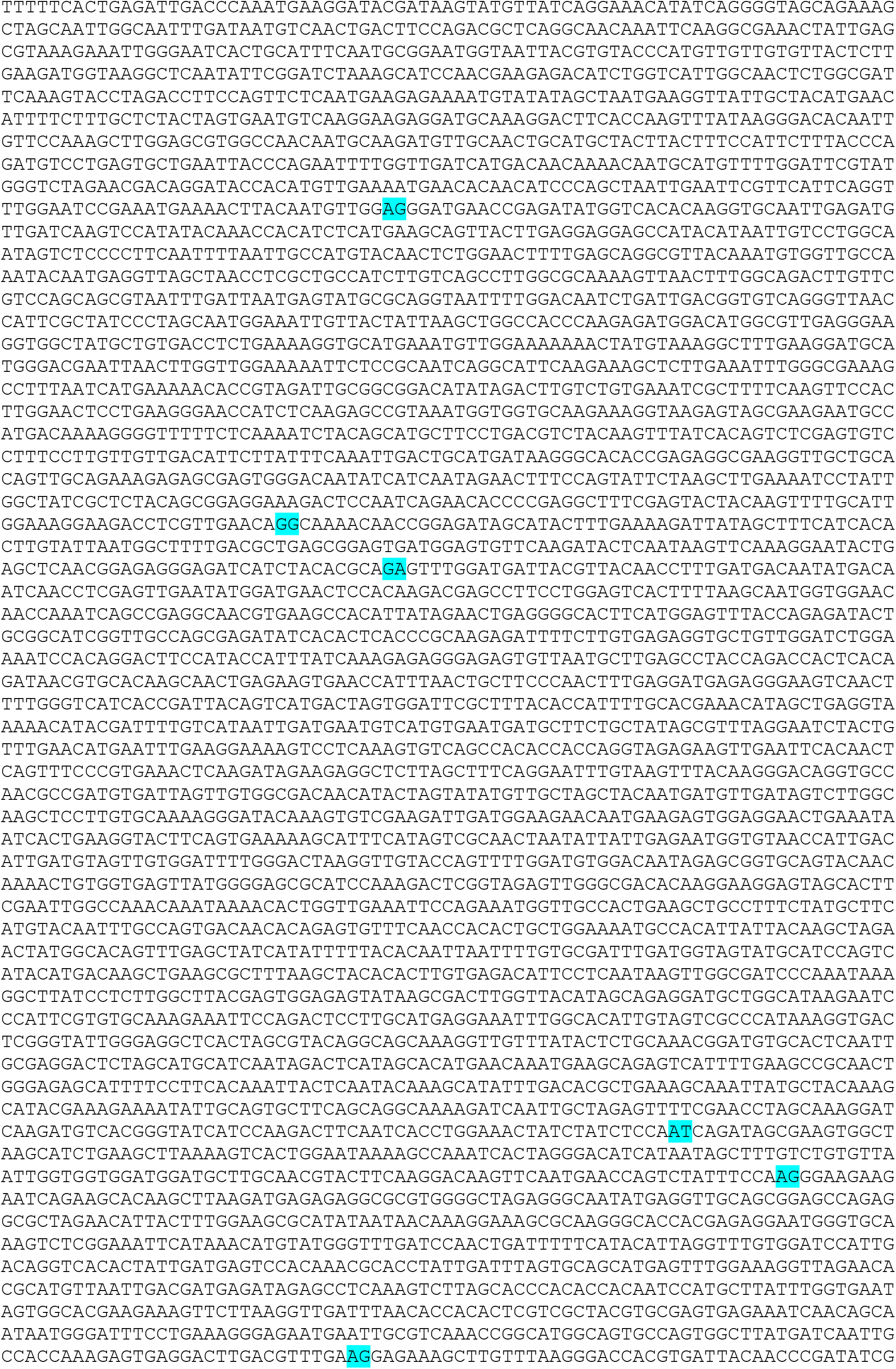
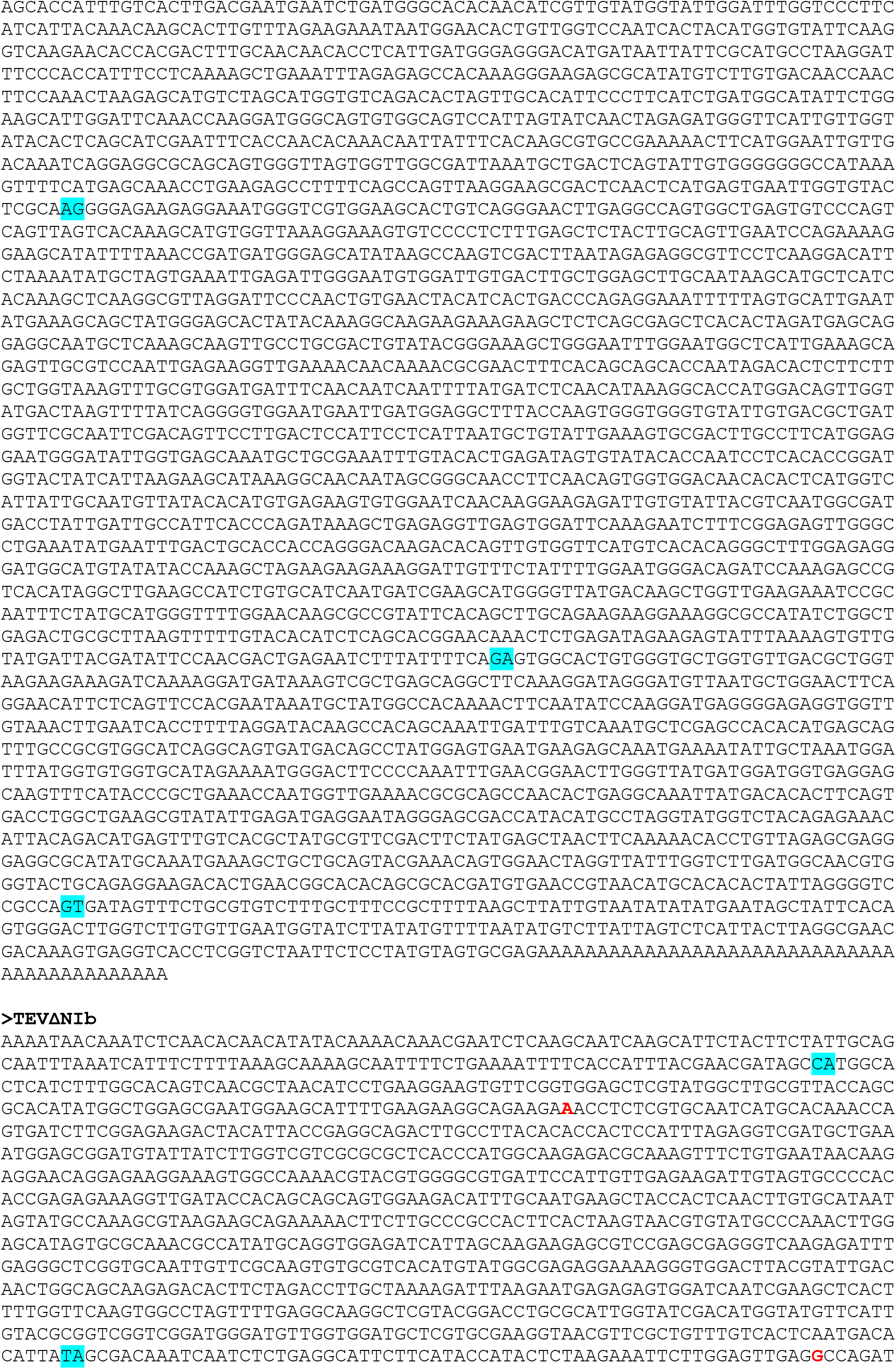
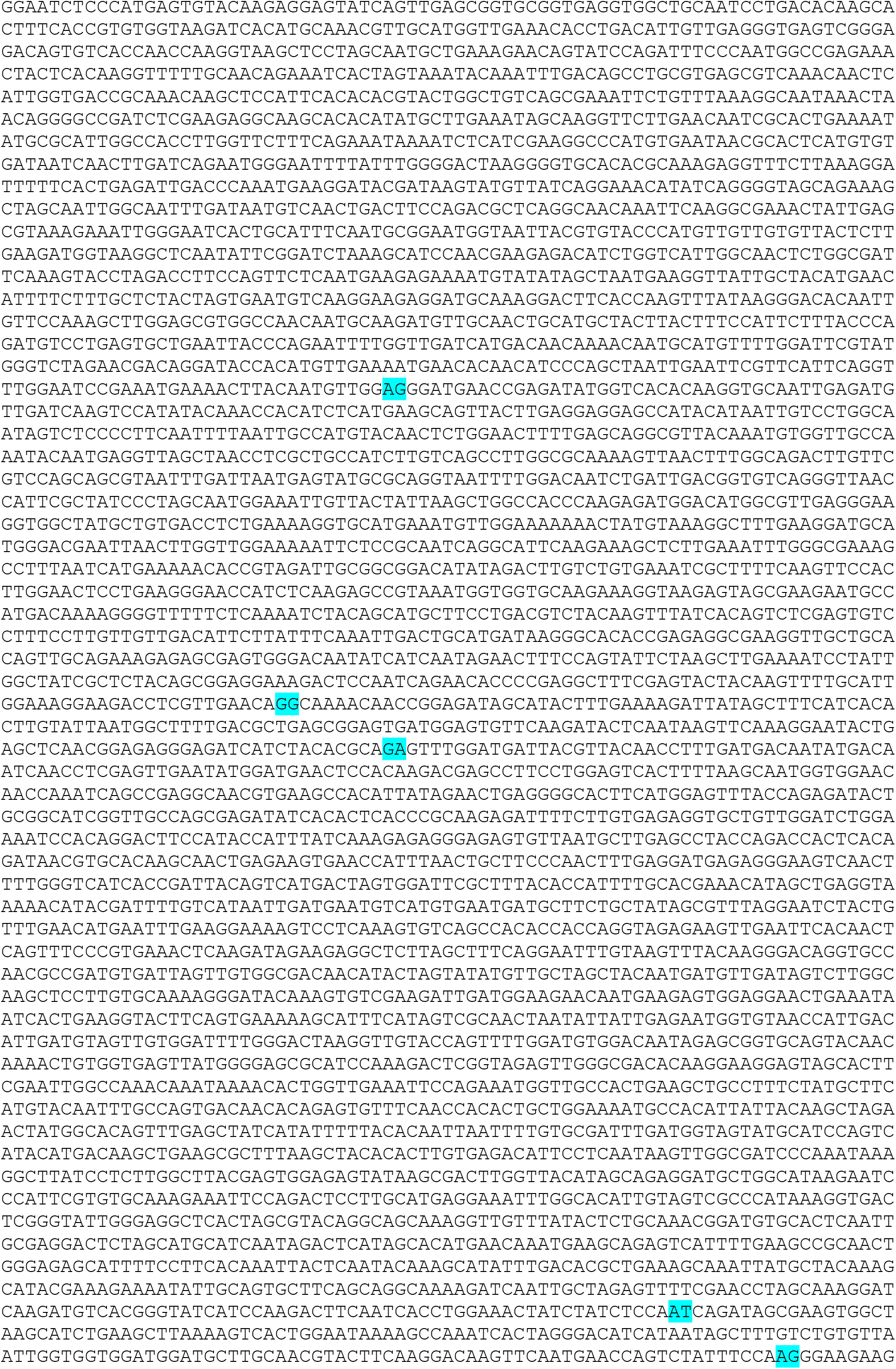
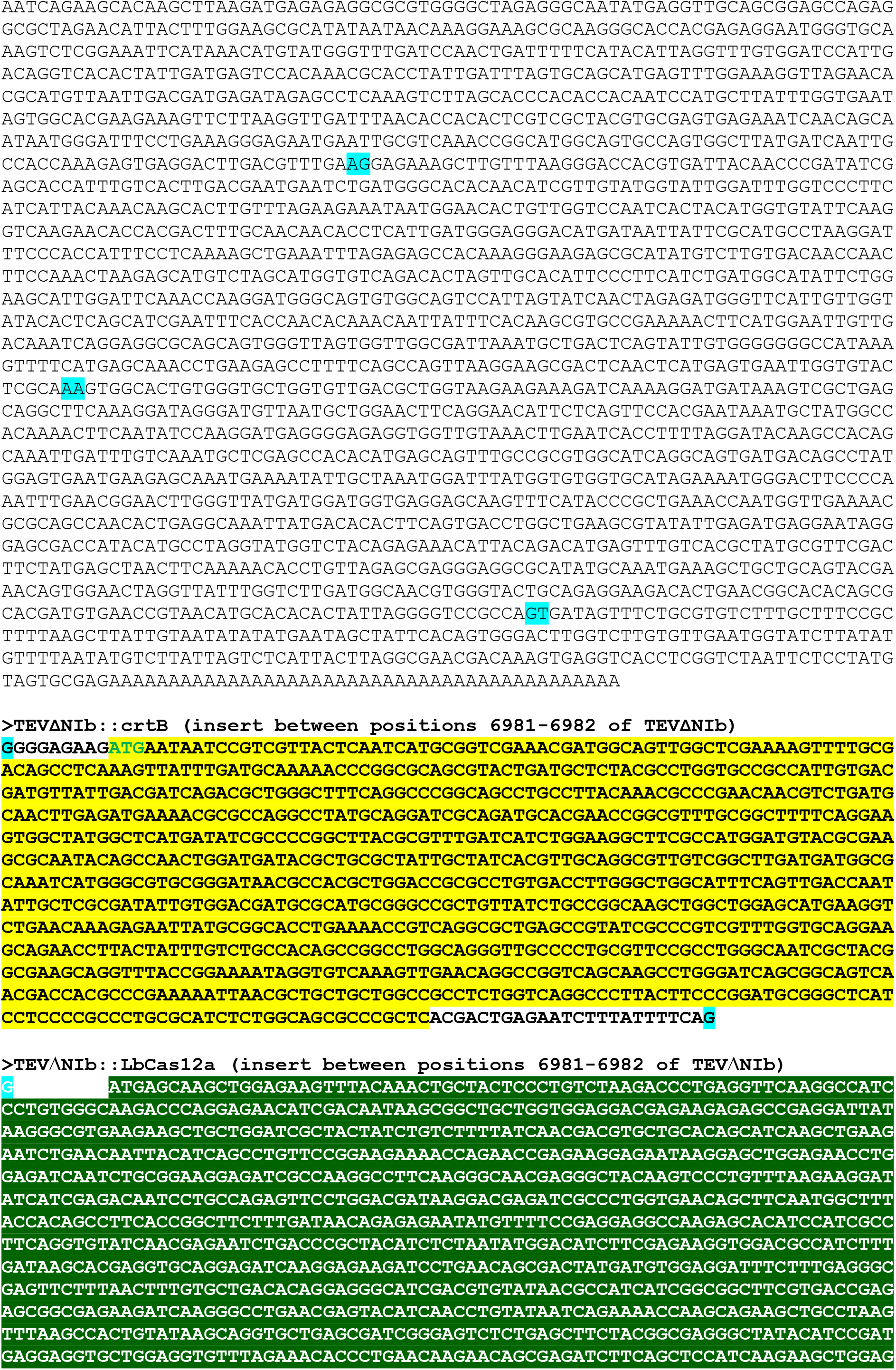
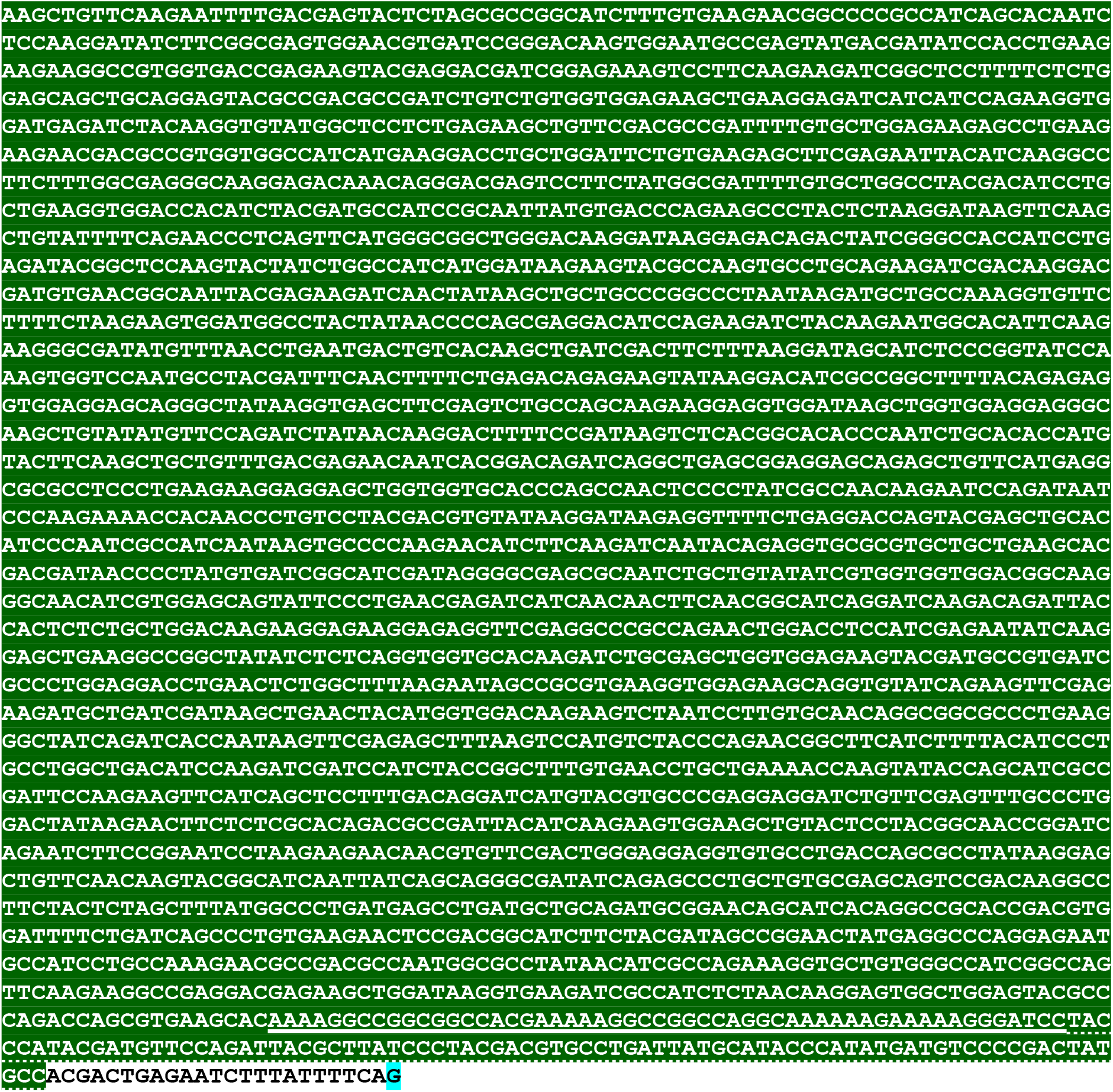
Full sequence of the wild-type tobacco etch virus (TEV), the defective viral vector TEVΔNIb and their derived recombinant viruses, TEVΔNIb::crtB and TEVΔNIb::LbCas12a. TEV-wt sequence corresponds to Genbank accession number DQ986288 including two silent mutations (G273A and A1119G, in red). Limits between TEV cistrons are marked on blue background. cDNAs corresponding to *P. ananatis* **phytoene synthase** (crtB) and Cas12a from ***Lachnospiraceae bacterium*** **ND2006** (LbCas12a) are on yellow and dark green backgrounds, respectively. <u>Nucleoplasmin nuclear localization signal (NLS)</u> and <u>human influenza hemagglutinin (3xHA) tag</u> are underlined and dotted, respectively. In the inserted cDNAs, sequences corresponding to **native** and **artificial** TEV NIaPro cleavage sites are in black or blue, respectively.

**Figure S2.**
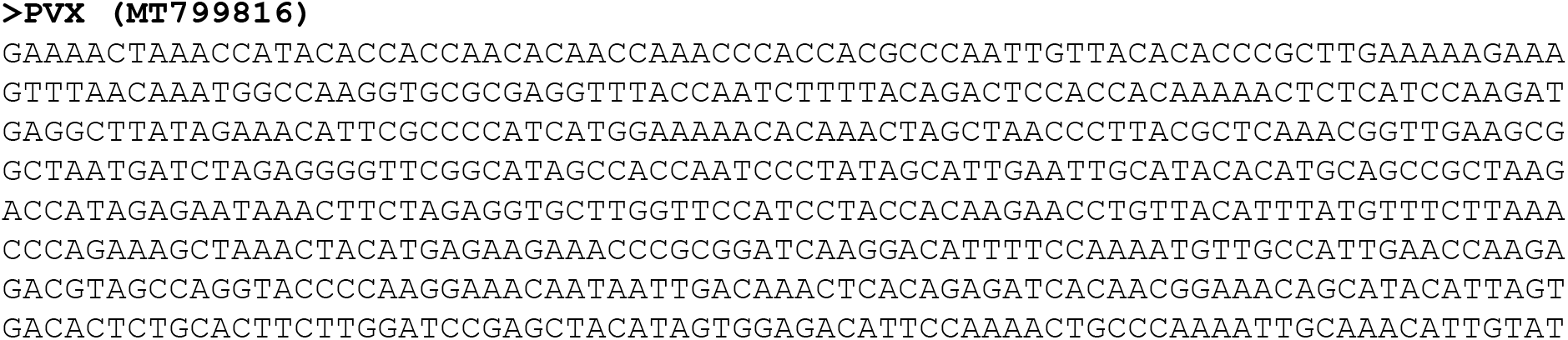
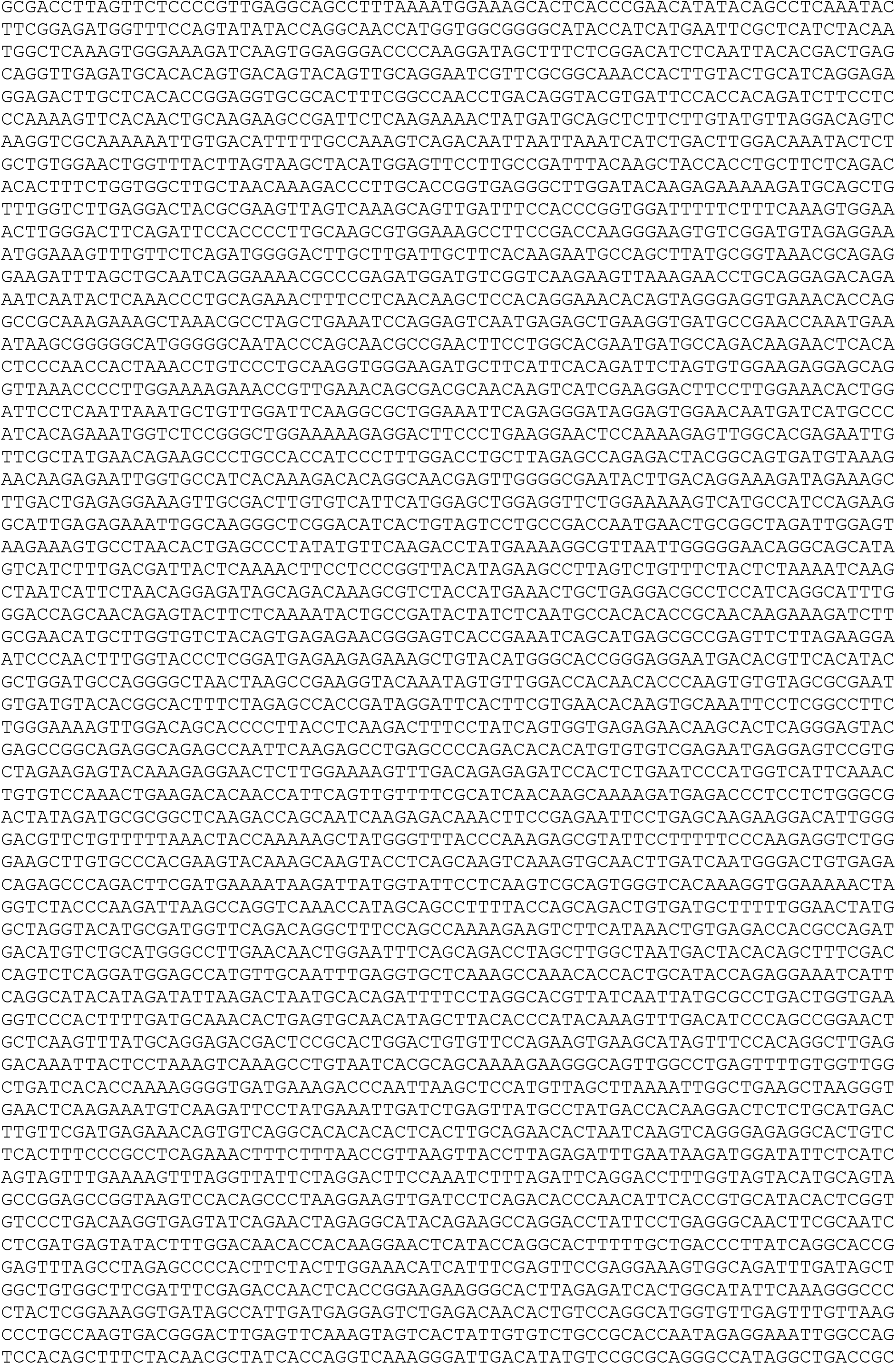
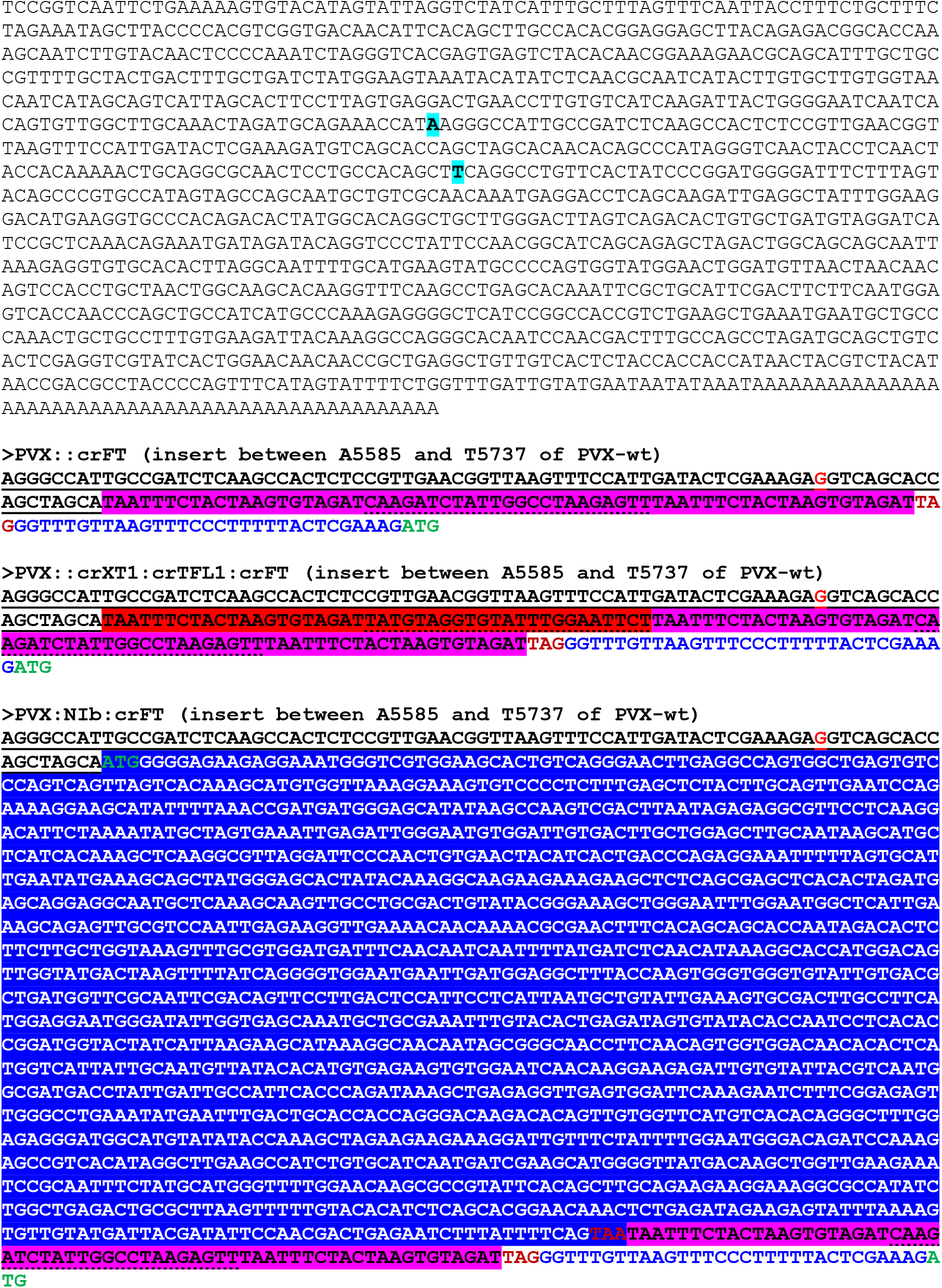
Full sequence of wild-type potato virus X (PVX; GenBank accession number MT799816) and its derived recombinant viruses PVX::crFT and PVX::NIb:crFT. In recombinant PVX clones, heterologous sequences are transcribed from coat protein (CP) promoter and a deleted version of PVX CP, lacking the 29 initial codons, is transcribed from an heterologous promoter derived from *Bamboo mosaic virus* (BaMV) CP (Dickmeis et al., 2014). **<u>PVX CP promoter</u>** with a ATG-AGG mutation to abolish start codon is underlined. **BaMV CP promoter** is in blue. crRNA sequences corresponding to ***NbFT*** and ***NbXT1*** are on pink and red background, respectively, and protospacer region is underlined (dotted). cDNA corresponding to **TEV nuclear inclusion *b* (NIb) cistron** is on dark blue background.

**Figure S3.**
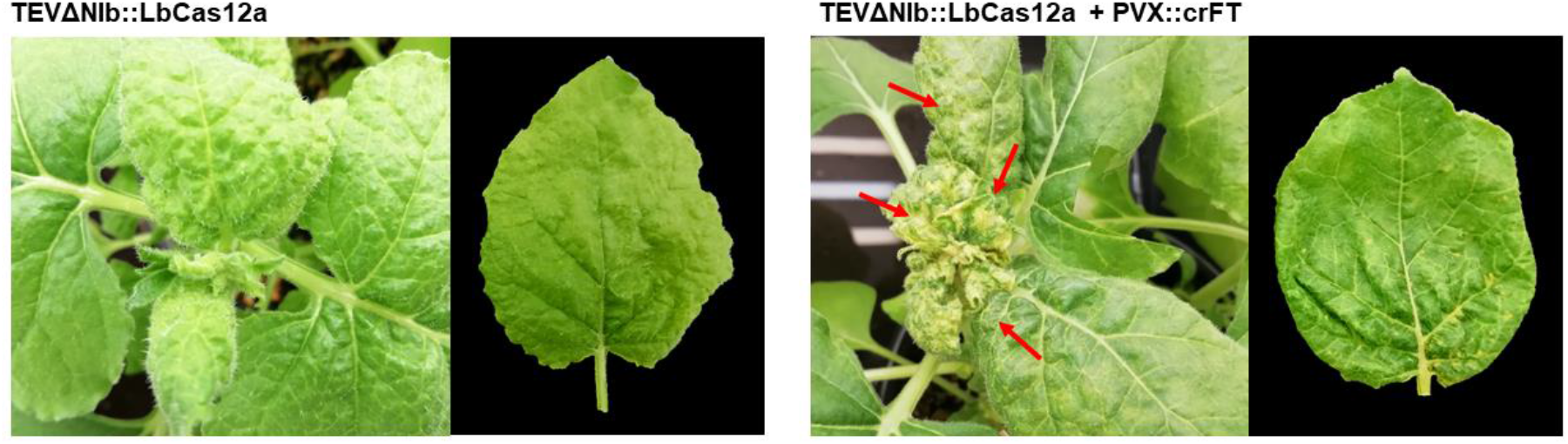
35S::NIb *N. benthamiana* plants and representative leaves from these plants at 14 dpi inoculated with TEVΔNIb::LbCas12a (left) or co-inoculated with TEVΔNIb::LbCas12a and PVX::crFT (right). Chlorotic spots in leaves from co-inoculated plants are indicated with red arrows.

**Figure S4.**
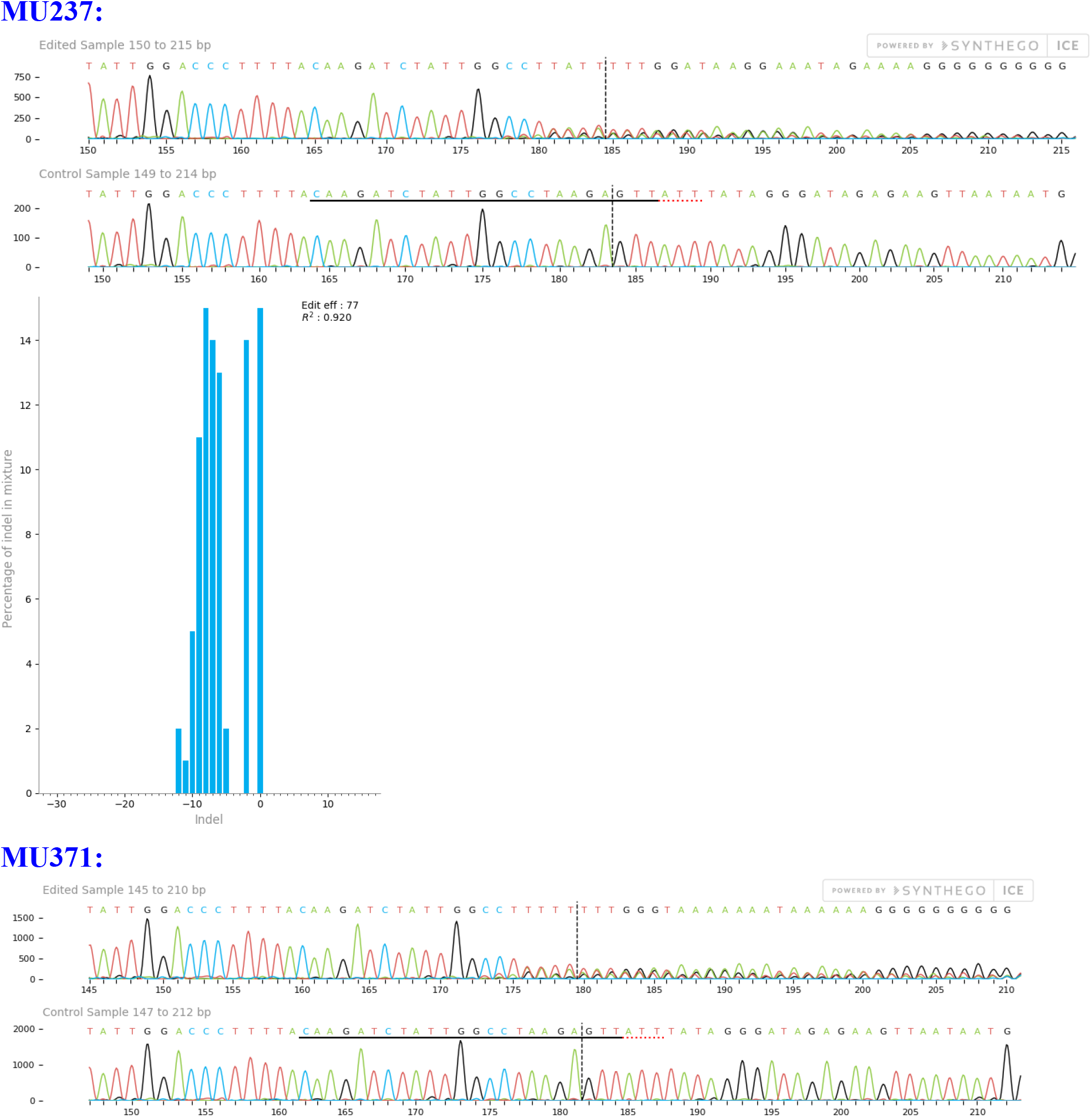
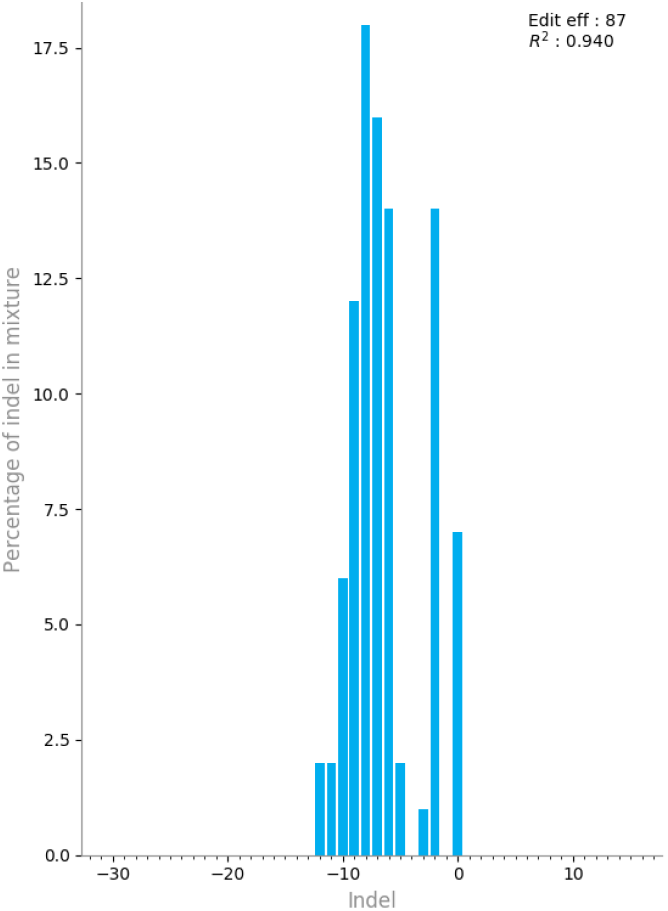
ICE analysis results of target site mutations in 35S::NIb *N. benthamiana* plants co-inoculated with TEVΔNIb::LbCas12a and PVX::crFT. Examples of sequencing chromatograms (top) and mutagenesis profiles (bottom) are shown.

**Figure S5.**
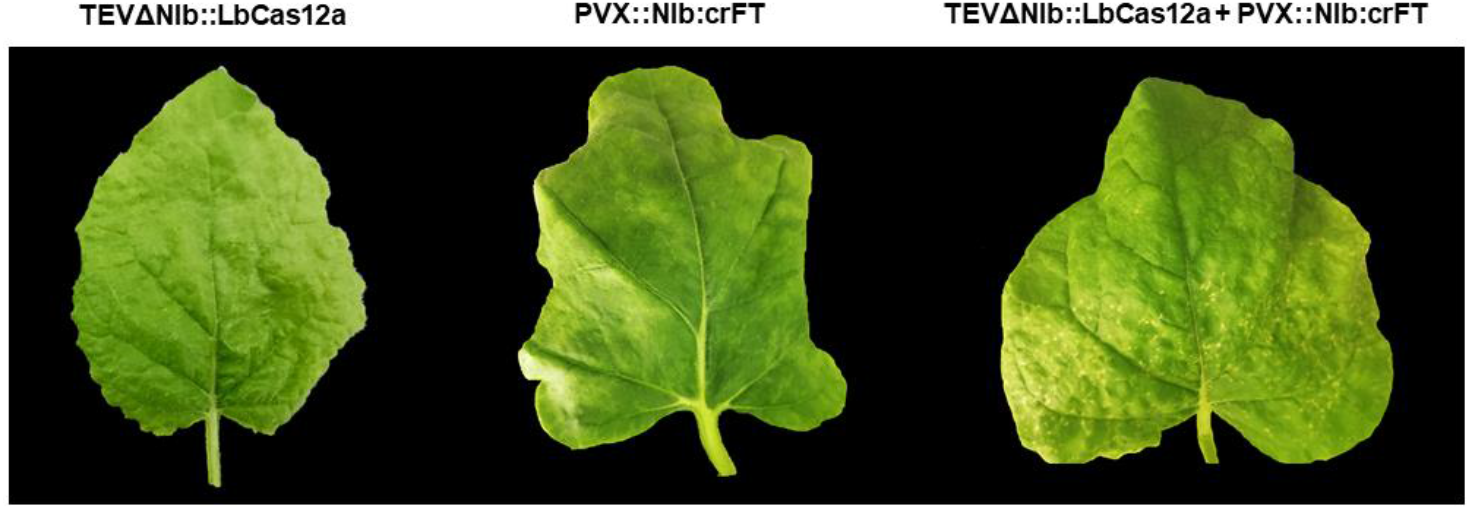
Representative leaves from wild-type *N. benthamiana* plants at 14 dpi inoculated with TEVΔNIb::LbCas12a alone (left), PVX::NIb:crFT alone (middle), or coinoculated with TEVΔNIb::LbCas12a and PVX::NIb:crFT (right).

**Table S1.**
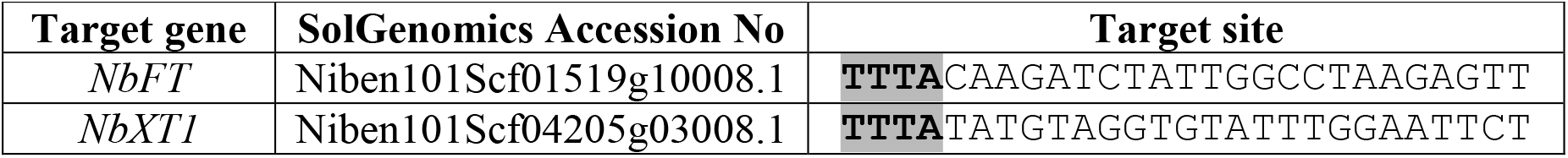
*N. benthamiana* genes targeted by the CRISPR-Cas12a system. Note that PAM sequence recognised by LbCas12a nuclease is highlighted in grey.

**Table S2.**
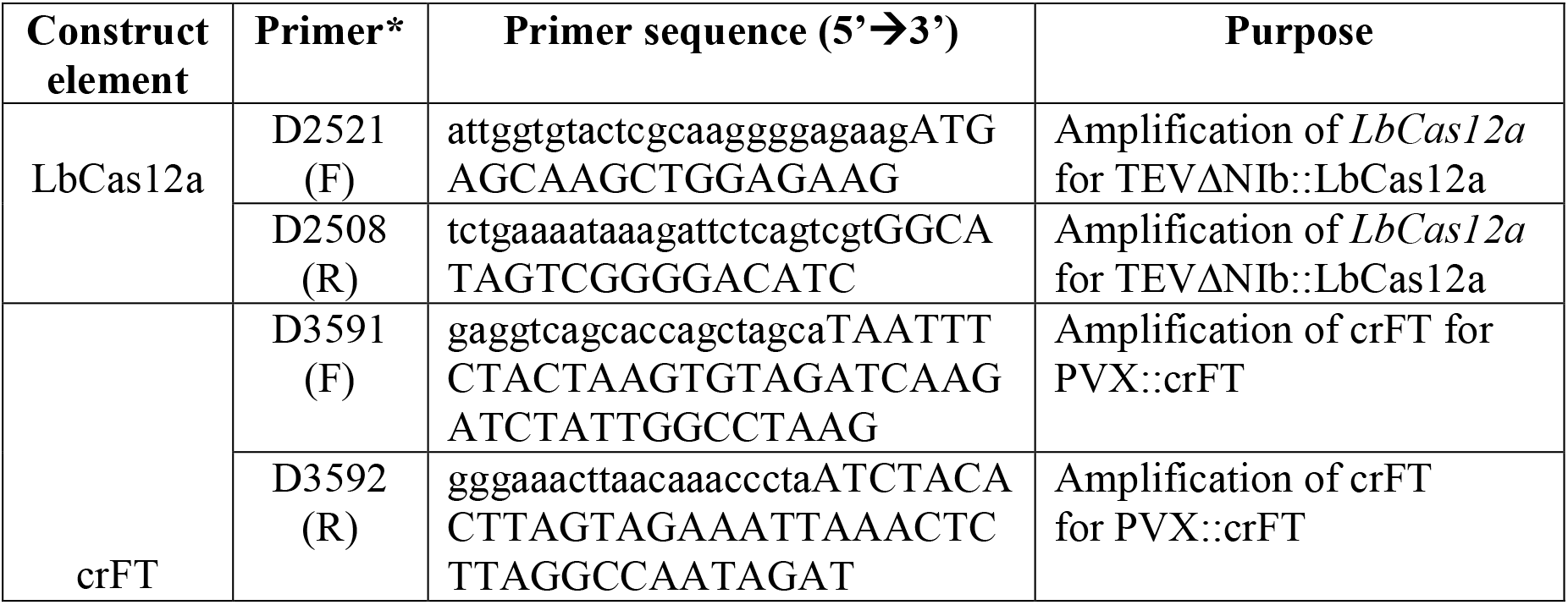
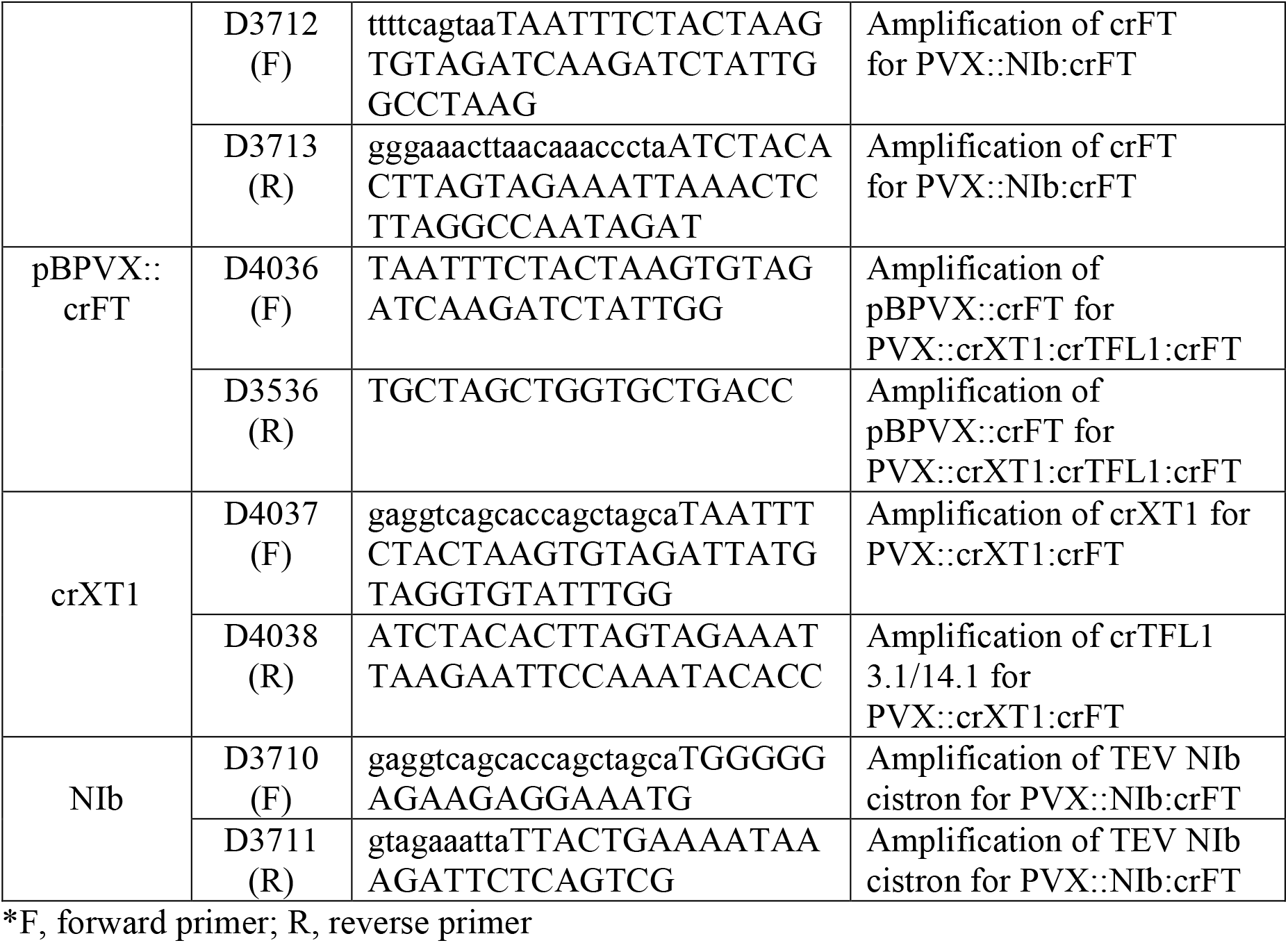
Primers used for the construction of recombinant viruses.

**Table S3.** Primer combinations used for the construction of recombinant viruses.

| Construct | Primer names |
| --- | --- |
| TEVΔNlb::LbCas12a | D2521 + D2508 (template: pcDNA3.1::hLbCpf1) |
| PVX::crFT | D3591+ D3592 (no template) |
| PVX::crXT1:crFT | D4037 + D4038<br>D4036 + D3536 (template: pBPVX::crFT) |
| PVX::Nlb:crFT | D3710 + D3711 (template: pGTEVa)<br>D3712 + D3713 |

**Table S4.**
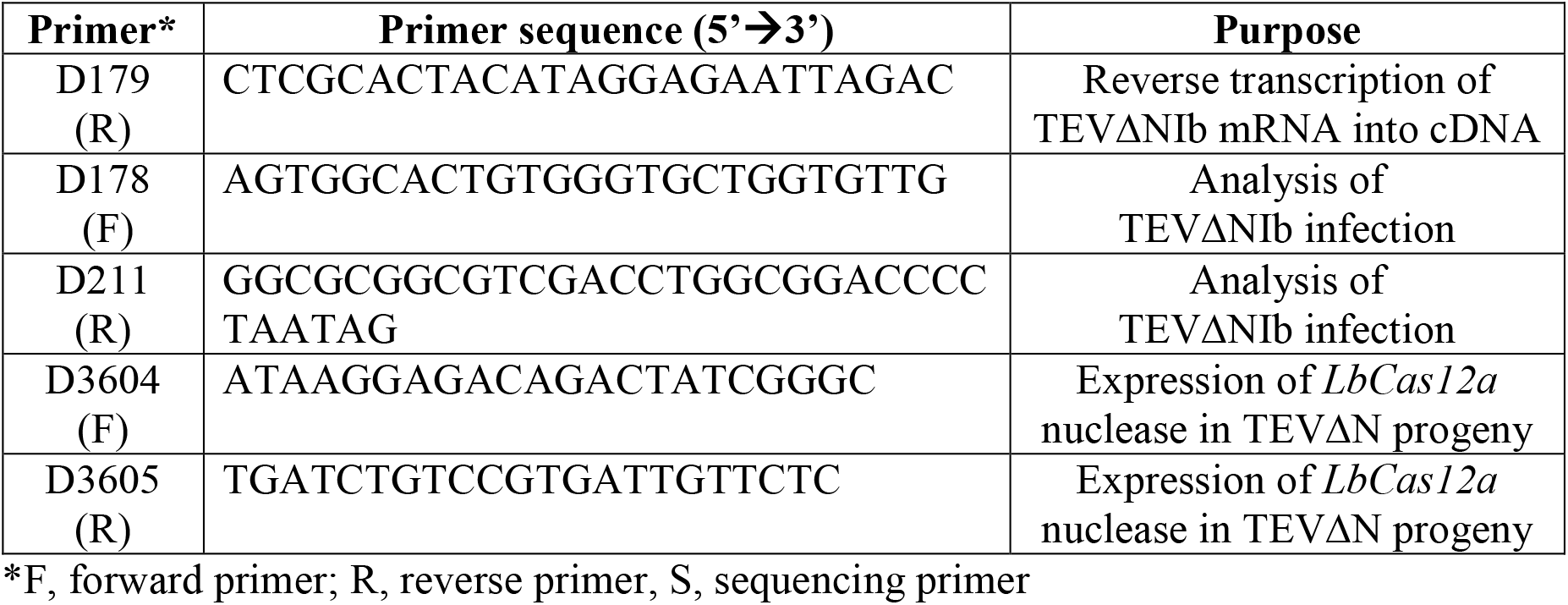
Primers used for TEVΔNIb diagnosis by RT-PCR.

**Table S5.**
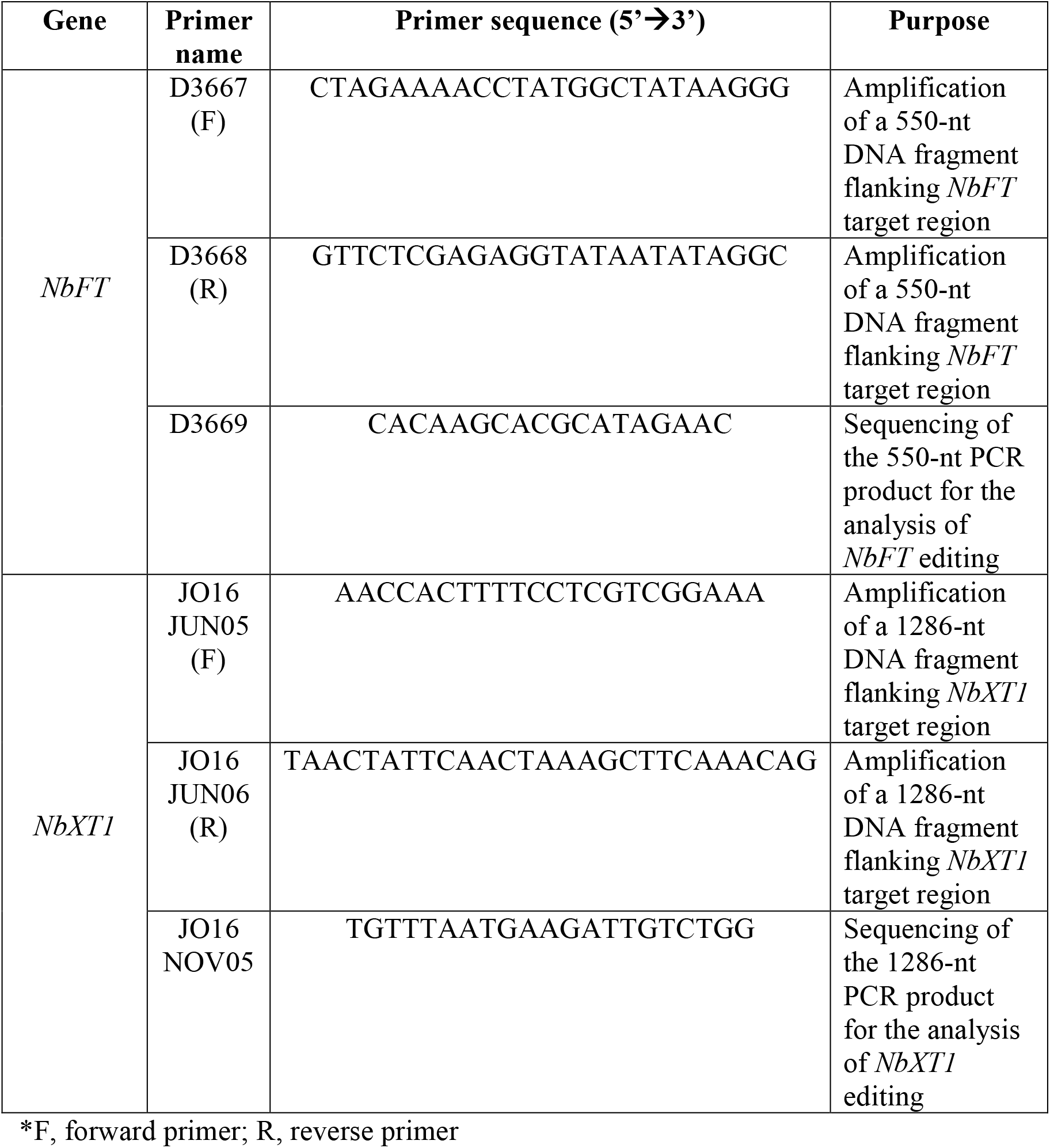
Primers used for Cas12a-crRNA gene editing analysis.

## References

1. Ali, Z., Eid, A., Ali, S., and Mahfouz, M.M. (2018) Pea early-browning virus-mediated genome editing via the CRISPR/Cas9 system in Nicotiana benthamiana and Arabidopsis. Virus Res., 244, 333–337.

2. Ariga, H., Toki, S., and Ishibashi, K. (2020) Potato virus X vector-mediated DNA-free genome editing in plants. Plant Cell Physiol., 61, 1946–1953.

3. Bedoya, L., Martínez, F., Rubio, L., and Daròs, J.A. (2010) Simultaneous equimolar expression of multiple proteins in plants from a disarmed potyvirus vector. J. Biotechnol., 150, 268–275.

4. Bernabé-Orts, J.M., Casas-Rodrigo, I., Minguet, E.G., Landolfi, V., Garcia-Carpintero, V., Gianoglio, S., et al. (2019) Assessment of Cas12a-mediated gene editing efficiency in plants. Plant Biotechnol. J., 17, 1971–1984.

5. Cody, W.B., Scholthof, H.B., and Mirkov, T.E. (2017) Multiplexed gene editing and protein overexpression using a tobacco mosaic virus viral vector. Plant Physiol., 175, 23–35.

6. Cong, L., Ran, F.A., Cox, D., Lin, S., Barretto, R., Habib, N., et al. (2013) Multiplex genome engineering using CRISPR/Cas systems. Science (80-.)., 339, 819–823.

7. Dickmeis, C., Fischer, R., and Commandeur, U. (2014) Potato virus X-based expression vectors are stabilized for long-term production of proteins and larger inserts. Biotechnol. J., 9, 1369–1379.

8. Ellison, E.E., Nagalakshmi, U., Gamo, M.E., Huang, P. jui, Dinesh-Kumar, S., and Voytas, D.F. (2020) Multiplexed heritable gene editing using RNA viruses and mobile single guide RNAs. Nat. Plants, 6, 620–624.

9. Endo, A., Masafumi, M., Kaya, H., and Toki, S. (2016) Efficient targeted mutagenesis of rice and tobacco genomes using Cpf1 from Francisella novicida. Sci. Rep., 6, 38169.

10. Gilbertson, R.L., Sudarshana, M., Jiang, H., Rojas, M.R., and Lucas, W.J. (2003) Limitations on Geminivirus Genome Size Imposed by Plasmodesmata and Virus-Encoded Movement Protein: Insights into DNA Trafficking. Plant Cell, 15, 2578–2591.

11. Hu, J., Li, S., Li, Z., Li, H., Song, W., Zhao, H., et al. (2019) A barley stripe mosaic virus-based guide RNA delivery system for targeted mutagenesis in wheat and maize. Mol. Plant Pathol., 20, 1463–1474.

12. Huang, T.K. and Puchta, H. (2021) Novel CRISPR/Cas applications in plants: from prime editing to chromosome engineering. Transgenic Res.

13. Jackson, A.O. and Christie, S.R. (1977) Purification and some physicochemical properties of sonchus yellow net virus. Virology, 77, 344–355.

14. Jiang, N., Zhang, C., Liu, J.Y., Guo, Z.H., Zhang, Z.Y., Han, C.G., and Wang, Y. (2019) Development of Beet necrotic yellow vein virus-based vectors for multiple-gene expression and guide RNA delivery in plant genome editing. Plant Biotechnol. J., 17, 1302–1315.

15. Kim, D., Kim, J., Hur, J.K., Been, K.W., Yoon, S.-H., and Kim, J.-S. (2016) Genome-wide analysis reveals specificities of Cpf1 endonucleases in human cells. Nat. Biotechnol., 34, 863–8.

16. Kim, H., Kim, S.T., Ryu, J., Kang, B.C., Kim, J.S., and Kim, S.G. (2017) CRISPR/Cpf1-mediated DNA-free plant genome editing. Nat. Commun., 8, 14406.

17. Kim, H.K., Song, M., Lee, J., Menon, A.V., Jung, S., Kang, Y.M., et al. (2017) In vivo high-throughput profiling of CRISPR-Cpf1 activity. Nat. Methods, 14, 153–159.

18. Lau, C.H. and Suh, Y. (2017) In vivo genome editing in animals using AAV-CRISPR system: Applications to translational research of human disease. F1000Research, 6, 2153.

19. Lei, J., Dai, P., Li, Y., Zhang, W., Zhou, G., Liu, C., and Liu, X. (2021) Heritable gene editing using FT mobile guide RNAs and DNA viruses. Plant Methods, 17.

20. Li, J., Zhang, H., Si, X., Tian, Y., Chen, K., Liu, J., et al. (2017) Generation of thermosensitive male-sterile maize by targeted knockout of the ZmTMS5 gene. J. Genet. Genomics, 44, 465–468.

21. Loebenstein, G. and Gaba, V. (2012) Viruses of Potato. In: Advances in Virus Research, pp. 209–246. Academic Press Inc.

22. Ma, X., Zhang, X., Liu, H., and Li, Z. (2020) Highly efficient DNA-free plant genome editing using virally delivered CRISPR–Cas9. Nat. Plants, 6, 773–779.

23. Majer, E., Llorente, B., Rodríguez-Concepción, M., and Daròs, J.A. (2017) Rewiring carotenoid biosynthesis in plants using a viral vector. Sci. Rep., 7, 41645.

24. Martí, M., Diretto, G., Aragonés, V., Frusciante, S., Ahrazem, O., Gómez-Gómez, L., and Daròs, J.A. (2020) Efficient production of saffron crocins and picrocrocin in Nicotiana benthamiana using a virus-driven system. Metab. Eng., 61, 238–250.

25. Nekrasov, V., Staskawicz, B., Weigel, D., Jones, J.D.G., and Kamoun, S. (2013) Targeted mutagenesis in the model plant Nicotiana benthamiana using Cas9 RNA-guided endonuclease. Nat. Biotechnol., 31, 691–693.

26. Peng, X., Ma, X., Lu, S., and Li, Z. (2021) A Versatile Plant Rhabdovirus-Based Vector for Gene Silencing, miRNA Expression and Depletion, and Antibody Production. Front. Plant Sci., 11, 627880.

27. Platt, R.J., Chen, S., Zhou, Y., Yim, M.J., Swiech, L., Kempton, H.R., et al. (2014) CRISPR-Cas9 knockin mice for genome editing and cancer modeling. Cell, 159, 440–455.

28. Senís, E., Fatouros, C., Große, S., Wiedtke, E., Niopek, D., Mueller, A.-K., et al. (2014) CRISPR/Cas9-mediated genome engineering: an adeno-associated viral (AAV) vector toolbox. Biotechnol. J., 9, 1402–12.

29. Tang, X., Lowder, L.G., Zhang, T., Malzahn, A.A., Zheng, X., Voytas, D.F., et al. (2017) A CRISPR-Cpf1 system for efficient genome editing and transcriptional repression in plants. Nat. Plants, 3, 17103.

30. Uranga, M., Aragonés, V., Selma, S., Vázquez-Vilar, M., Orzáez, D., and Daròs, J. (2021) Efficient Cas9 multiplex editing using unspaced sgRNA arrays engineering in a Potato virus X vector. Plant J.

31. Wang, M., Mao, Y., Lu, Y., Tao, X., and Zhu, J.-K. (2017) Multiplex Gene Editing in Rice Using the CRISPR-Cpf1 System. Mol. Plant, 10, 1011–1013.

32. Wylie, S.J., Adams, M., Chalam, C., Kreuze, J., López-Moya, J.J., Ohshima, K., et al. (2017) ICTV virus taxonomy profile: Potyviridae. J. Gen. Virol., 98, 352–354.

33. Xu, C.L., Ruan, M.Z.C., Mahajan, V.B., and Tsang, S.H. (2019) Viral delivery systems for CRISPR. Viruses, 11, 28.

34. Xu, R., Qin, R., Li, H., Li, D., Li, L., Wei, P., and Yang, J. (2017) Generation of targeted mutant rice using a CRISPR-Cpf1 system. Plant Biotechnol. J., 15, 713–717.

35. Yin, K., Han, T., Liu, G., Chen, T., Wang, Y., Yu, A.Y.L., and Liu, Y. (2015) A geminivirus-based guide RNA delivery system for CRISPR/Cas9 mediated plant genome editing. Sci. Rep., 5, 14926.

36. Zahir Ali, Aala Abul-faraj, Lixin Li, Neha Ghosh, Marek Piatek, Ali Mahjoub, Mustapha Aouida, Agnieszka Piatek, Nicholas J Baltes, Daniel F Voytas, Savithramma Dinesh-Kumar, M.M.M. (2015) Efficient Virus-Mediated Genome Editing in Plants Using the CRISPR/Cas9 System. Mol. Plant, 8, 1288–91.

37. Zaidi, S.S.-A., Magdy M. Mahfouz, A., and Mansoor, S. (2017) CRISPR-Cpf1: A New Tool for Plant Genome Editing. Trends Plant Sci., 22, 549–550.

38. Zaidi, S.S. e. A., Mahfouz, M.M., and Mansoor, S. (2017) CRISPR-Cpf1: A New Tool for Plant Genome Editing. Trends Plant Sci., 22, 550–553.

39. Zetsche, B., Gootenberg, J.S., Abudayyeh, O.O., Slaymaker, I.M., Makarova, K.S., Essletzbichler, P., et al. (2015) Cpf1 Is a Single RNA-Guided Endonuclease of a Class 2 CRISPR-Cas System. Cell, 163, 759–771.

40. Zetsche, B., Heidenreich, M., Mohanraju, P., Fedorova, I., Kneppers, J., Degennaro, E.M., et al. (2017) Multiplex gene editing by CRISPR-Cpf1 using a single crRNA array. Nat. Biotechnol., 35, 31–34.

41. Zhang, Y., Liang, Z., Zong, Y., Wang, Y., Liu, J., Chen, K., et al. (2016) Efficient and transgene-free genome editing in wheat through transient expression of CRISPR/Cas9 DNA or RNA. Nat. Commun., 7, 12617.

42. Zhu, H., Li, C., and Gao, C. (2020) Applications of CRISPR–Cas in agriculture and plant biotechnology. Nat. Rev. Mol. Cell Biol., 21, 661–677.

